# Multimodal Surface Matching with Higher-Order Smoothness Constraints^☆^

**DOI:** 10.1101/178962

**Authors:** Emma C. Robinson, Kara Garcia, Matthew F. Glasser, Zhengdao Chen, Timothy S. Coalson, Antonios Makropoulos, Jelena Bozek, Robert Wright, Andreas Schuh, Matthew Webster, Jana Hutter, Anthony Price, Lucilio Cordero Grande, Emer Hughes, Nora Tusor, Philip V. Bayly, David C. Van Essen, Stephen M. Smith, A. David Edwards, Joseph Hajnal, Mark Jenkinson, Ben Glocker, Daniel Rueckert

**Author notes:** Corresponding author Email address (Emma C. Robinson). These authors contributed equally (Kara Garcia, Matthew F. Glasser).

## Abstract

In brain imaging, accurate alignment of cortical surfaces is fundamental to the statistical sensitivity and spatial localisation of group studies; and cortical surface-based alignment has generally been accepted to be superior to volume-based approaches at aligning cortical areas. However, human subjects have considerable variation in cortical folding, and in the location of functional areas relative to these folds. This makes alignment of cortical areas a challenging problem. The Multimodal Surface Matching (MSM) tool is a flexible, spherical registration approach that enables accurate registration of surfaces based on a variety of different features. Using MSM, we have previously shown that driving cross-subject surface alignment, using areal features, such as resting state-networks and myelin maps, improves group task fMRI statistics and map sharpness. However, the initial implementation of MSM's regularisation function did not penalize all forms of surface distortion evenly. In some cases, this allowed peak distortions to exceed neurobiologically plausible limits, unless regularisation strength was increased to a level which prevented the algorithm from fully maximizing surface alignment. Here we propose and implement a new regularisation penalty, derived from physically relevant equations of strain (deformation) energy, and demonstrate that its use leads to improved and more robust alignment of multimodal imaging data. In addition, since spherical warps incorporate projection distortions that are unavoidable when mapping from a convoluted cortical surface to the sphere, we also propose constraints that enforce smooth deformation of cortical anatomies. We test the impact of this approach for longitudinal modelling of cortical development for neonates (born between 32 and 45 weeks of post-menstrual age) and demonstrate that the proposed method increases the biological interpretability of the distortion fields and improves the statistical significance of population-based analysis relative to other spherical methods.

## 1. Introduction

The cerebral cortex is a highly convoluted sheet, with complex patterns of folding that vary considerably across individuals. Accurate cross-subject volumetric registration, in the face of folding variability, is far from straightforward as relatively small deformations in three dimensions (3D) risk matching opposing banks of cortical folds, or aligning brain tissue with cerebrospinal fluid. For this reason surface registration methods have been proposed, which constrain alignment to the 2D cortical sheet (Durrleman et al., 2009; Fischl et al., 1999b; Gu et al., 2004; Lombaert et al., 2013; Lyu et al., 2015; Robinson et al., 2014; Tsui et al., 2013; Wright et al., 2015; Yeo et al., 2010; Van Essen, 2005). These are generally accepted to have better performance at aligning cortical areas (Glasser et al., 2016b)

Often, surface registration techniques have focused on the alignment of cortical convolutions. Examples include, spectral embedding approaches, which learn fast and accurate mappings between low-dimensional representations of cortical shapes (Lombaert et al., 2013; Orasanu et al., 2016b; Wright et al., 2015); Large Deformation Diffeomorphic Metric Mapping (LDDMM) frameworks, which learn vector flows fields between cortical geometries (Durrleman et al., 2009), to allow smooth deformation of cortical shapes (Durrleman et al., 2013); and spherical projection methods (Fischl et al., 1999b; Lyu et al., 2015; Van Essen et al., 2012; Yeo et al., 2010), which simplify the problem of cortical registration by projecting the convoluted surface to a sphere. All methods demonstrate clear advantages in terms of: improving the speed and accuracy of alignment (Lombaert et al., 2013; Wright et al., 2015; Yeo et al., 2010); increasing the correspondence of important features on the cortical surface, such as brain activations (Fischl et al., 2008; Lombaert et al., 2015; Yeo et al., 2010); and providing a platform through which cortical shapes can be statistically compared (Durrleman et al., 2013; Orasanu et al., 2016b)

Ultimately, however, shape based alignment of the brain is limited as cortical folding patterns vary considerably across individuals, and correspond poorly with the placement of cortical architecture, function, connectivity, and topographic maps across most parts of the cortex (Amunts et al., 2000, 2007; Glasser et al., 2016a). For this reason several papers have been proposed to drive cortical alignment using ‘areal’ features (descriptors that correlate with the functional organisation of the human brain). These include Conroy et al. (2013); Frost and Goebel (2013); Nenning et al. (2017); Sabuncu et al. (2010), which drive spherical registration using features derived from functional Magnetic Resonance Imaging (fMRI), and Tardif et al. (2015), who register level-set representations of cortical volumes using combinations of geometric features and cortical myelin^2^. Further, in Lombaert et al. (2015) and Orasanu et al. (2016a) spectral shape embedding approaches are extended to utilise broader feature sets, with Lombaert et al. (2015) adapting spectral alignment of the visual cortex to improve transfer of retinotopic maps, and Orasanu et al. (2016a) improving correspondence matching between neonatal feature sets by performing multimodal spectral embeddings of cortical shape and diffusion MRI (dMRI).

Most of these methods are optimised for alignment of specialised feature sets, whether that be fMRI time series (Sabuncu et al., 2010), functional connectivity patterns (Conroy et al., 2013; Nenning et al., 2017) or fixed combinations of cortical geometry with fMRI/dMRI (Lombaert et al., 2015; Orasanu et al., 2016a). Furthermore, methods which enforce geometric constraints on the mapping (Durrleman et al., 2013; Lombaert et al., 2013, 2015; Orasanu et al., 2016b; Wright et al., 2015) are not well placed to address the known mismatch between the functional (areal) and morphological (folding) organisation of the human cerebral cortex (Amunts et al., 2000; Glasser et al., 2016a; Nenning et al., 2017)

Therefore, in Robinson et al. (2014) we proposed Multimodal Surface Matching (MSM), a spherical deformation approach that enables flexible alignment of any type or combination of features that can be mapped to the cortical surface. MSM is inspired by DROP (Glocker et al., 2008) a discrete optimisation approach for non-rigid registration of 3D volumes. Like DROP, MSM has advantages in terms of reduced sensitivity to local minima (Glocker et al., 2011), and a modular optimisation framework. By allowing any combination of similarity and regularisation terms, this framework can be used to align any type or combination of features, provided that an appropriate similarity cost can be found. Accordingly, MSM has been used to drive alignment of a wide variety of different feature sets, including cortical folding, cortical myelination, resting-state network maps, and multimodal combinations of folding and myelin (Bo΂ek et al., 2016; Glasser et al., 2016a,b; Harrison et al., 2015; Robinson et al., 2014). Experiments show that this also leads to improvements in correspondence of unrelated areal features including retinotopic maps (Abdollahi et al., 2014) and task fMRI (Glasser et al., 2016a; Robinson et al., 2014; Tavor et al., 2016).

One limitation of the original MSM implementation was that it only offered a first-order (pairwise) cost function to penalize against large distortions. Unfortunately, this was suboptimal, as the penalty for doubling the size of a region equalled that of reducing the area to zero. Further, it was not possible to adjust the weight between isotropic (size-changing) and anisotropic (shape-changing) distortions. We therefore required a method for considering higher-order interactions, where changes in both size and shape could be measured through triplets (triangles) of control-points. This was implemented in our discrete optimisation framework using clique reduction (Ishikawa, 2009, 2014), which has already been successfully applied to 2D registration in Glocker et al. (2010)

A prototype of this framework proved fundamental to development of the HCP’s multimodal parcellation (Glasser et al., 2016b,a). For Glasser et al. (2016a), we used a triplet-based Angular Deviation Penalty (ADP) that penalised change in angles for each triangular mesh face. Unfortunately, this new regularization penalty measure also had drawbacks. In particular, it did not have any direct penalty for increasing or decreasing the area of a feature. With careful tuning of the regularization strength, based on elimination of neurobiologically implausible individual subject peak distortions, we were able to produce the publicly released HCP results and the HCP’s multimodal parcellation (Glasser et al., 2016b,a). However, the need to eliminate neurobiologically implausible peak distortions limited the registration’s ability to maximize functional alignment. In particular, the method struggled to sufficiently penalize excessive distortion in regions where the individual topological layout of cortical areas deviated from the group (described in detail in Glasser et al. (2016a) Supplementary Methods sec 6.4). Similar evidence of topological variation has been reported in several other studies (Amunts et al., 2000; Gordon et al., 2017; Glasser et al., 2016a; Haxby et al., 2011; Wang et al., 2015).

For these reasons,in this paper we present higher-order MSM with a new penalty inspired by the hyperelastic properties of brain tissue. This mechanically-inspired penalty minimises surface deformations in a physically plausible way (Knutsen et al., 2010), fully controlling both size and shape distortions. We test this new strain-based regularization by re-optimizing the alignment of data from the adult Human Connectome Project (HCP), for both folding alignment (MSMSulc) and the multimodal alignment protocol (MSMAll) described in Glasser et al. (2016a) (sec 6.4). This uses myelin maps, resting state network maps, and resting state visuotopic maps to align cortical areas.

A further limitation of the original MSM framework has been that use of spherical alignment complicates the neurobiological interpretation of deformation fields. Specifically, spherical projection distorts the relative separation of vertices between the 3D anatomical surface (‘cortical anatomy’) and the sphere, such that spherical regularisation has a varying influence on different parts of the cortical anatomy (Fig. 1). These effects may vary across individuals or time points and depend on the projection algorithm used. Therefore, we propose an extension to the spherical alignment method, adapted from Knutsen et al. (2010), to deform points on the sphere but regularise displacements on the real anatomical surface mesh. This novel method retains the flexibility of the spherical framework (which allows registration of any type of feature that can be mapped to the cortical surface), while harnessing spatial information from the anatomical surface to produce physically appropriate warps.

**Figure 1.**
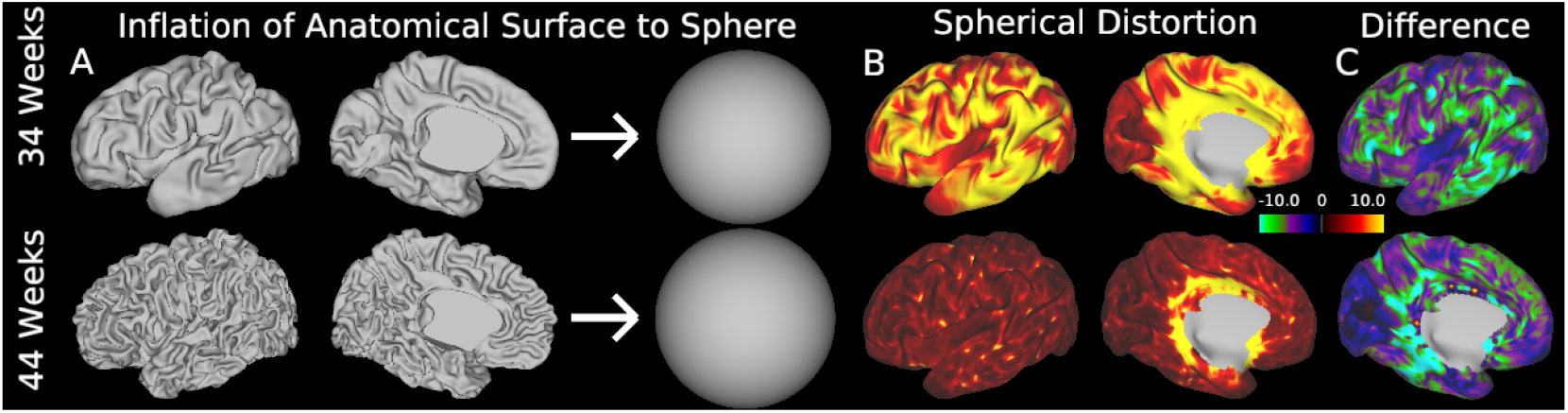
Areal distortions (changes in the relative spacing of vertices) occur as a result of projection from the anatomical surface to the spherical surface. These distortions change across the surfaces and between brains. Differences are particularly obvious for longitudinally acquired data. Shown here: A) White matter surfaces extracted from the same subject at 34 weeks post-menstrual age (PMA, top row) and 44 weeks PMA (bottom row) are projected to a sphere. B) Areal distortions estimated in terms of isotropic expansion of mesh faces (log_2_(Area_2_/Area_1_)), shown aligned and resampled to the 44 week subject (inflated brain view). C) Areal distortion difference between time points

We test the approach on alignment of 10 longitudinally-acquired neonatal cortical surface data sets imaged twice between 31 and 43 weeks post-menstrual age. By comparing the proposed method to other spherical alignment approaches, we explore whether the resulting deformation fields offer improved correspondence with expected growth trajectories over this developmental period.

The rest of paper is organised as follows: we first give an overview of the original spherical MSM method (sMSM), proposed in (Robinson et al., 2014) (sec. 2), and discuss the extension to higher-order regularisation penalty terms (sec. 3). Methods for approximating and regularising anatomical warps (aMSM) are then presented (sec. 4). Finally, experiments and results are presented on a feature sets derived from adult and developing Human Connectome Project (sec. 6). Code and configuration files for running these experiments are available (https://www.doc.ic.ac.uk/~ecr05/MSM_HŨCR_v2/). Preliminary dHCP data can be found from https://data.developingconnectome.org (version 1.1), and HCP data can be downloaded from https://db.humanconnectome.org^3^

## 2. Multimodal Surface Matching

We begin with an overview of the original MSM method, first proposed in Robinson et al. (2014). In this framework, we seek alignment between two anatomical surfaces, each projected to a sphere, through the procedures outlined in Fischl et al. (1999a). Here, anatomical surfaces represent tessellated meshes fit to the outer boundary of a white matter tissue segmentation (i.e. the gray/white surface). These are expanded outwards to the outer grey matter (or pial) boundary, and a midthickness surface is generated half way between white and pial boundaries. Separately, white matter surfaces are expanded to generate a smooth inflated cortical surface, from which points are then projected to a sphere. This is done in such a way as to minimise areal distortions i.e. minimise the overall change in area of triangular mesh faces during the transformation, (Fischl et al., 1999a). Further, during inflation, indexed vertices (or points) in each surface space retain one-to-one correspondence, such that each index represents the same cortical location across white, midthickness, pial, inflated and spherical surfaces.

The goal of MSM registration is to identify spatial correspondences between two spheres so as to improve the overlap of the surface geometry and/or functional properties of the cortical sheet from which the spheres were derived. Spheres may represent corresponding hemispheres (left or right) from: two different subjects, one subject and a population average template, or the same subject imaged at two different ages. During registration, vertex points on one (source) sphere are moved until the surface properties on that sphere better agree with those of the second (target) sphere. Due to the vertex correspondence between sphere and surface anatomy, this also implicitly derives correspondences for the cortical anatomy.

The registration process and surfaces involved are illustrated in Fig. 2. Let *SSS* be the source spherical surface with initial coordinates x, and *TSS* be the target spherical surface with coordinates **X**, where **x**, 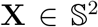. Let *SAS* be the anatomical (white/midthickness/pial) surface representation of *SSS* with coordinate y and TAS be the fixed anatomical surface representation of *TSS* with coordinates **Y**, where **y**, 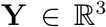. Let the moving source spherical surface be represented as *MSS* with coordinates x', and the resulting deformed anatomical configuration as (*DAS*) with coordinates y' (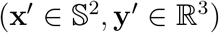). Anatomical surfaces may be white, midthickness, pial or inflated surfaces, and each surface is associated with multimodal feature sets **M** (fixed) and **m** (moving 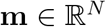). These represent any combination of *N* features describing cortical folding, brain function (such as resting state networks), cortical architecture, or structural connectivity.

**Figure 2.**
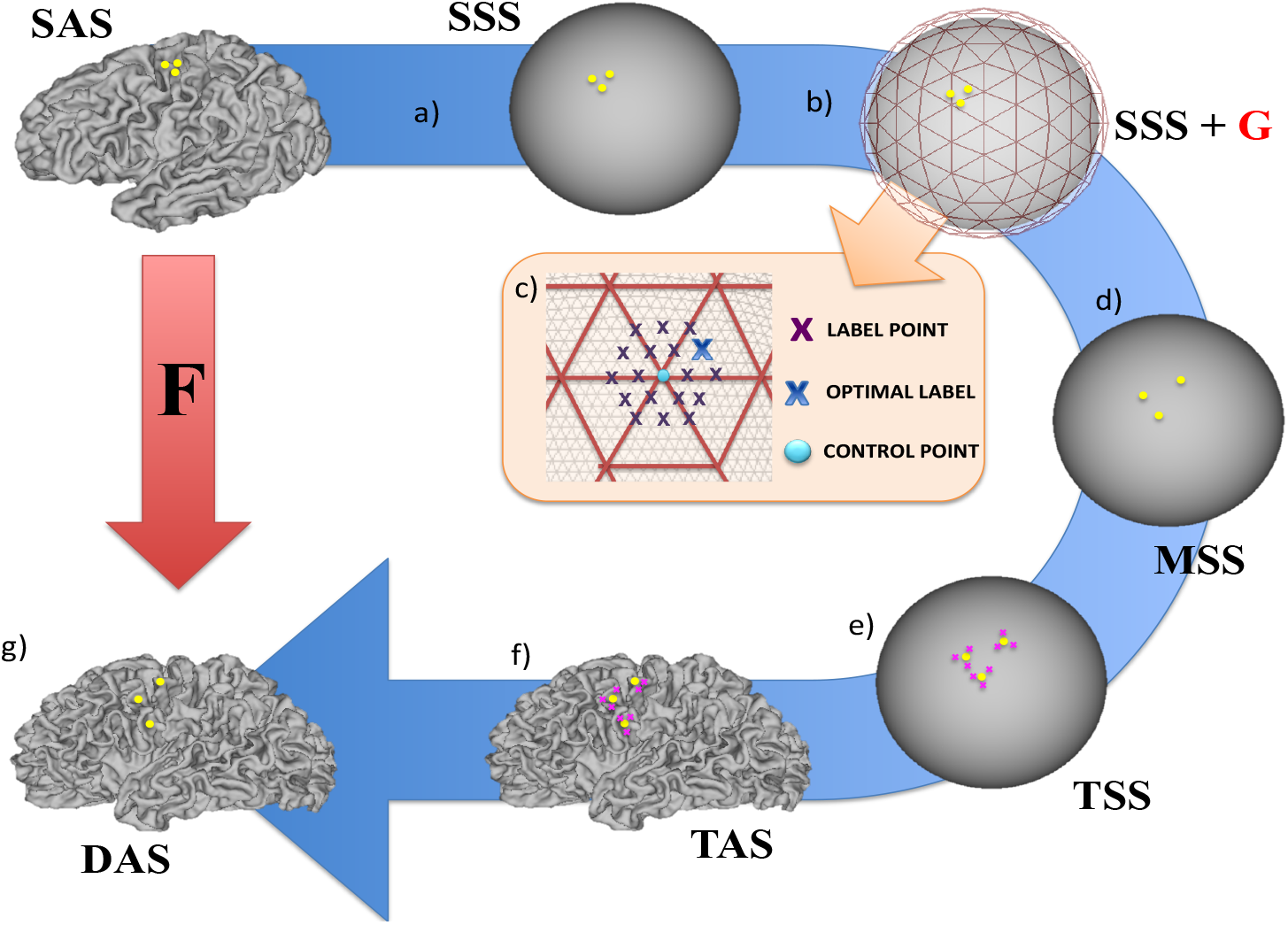
Projecting cortical anatomy through spherical warps. The figure follows the displacement of three (yellow) points on the source white anatomical surface (*SAS*), via the moving source spherical surface (*MSS*), into a new configuration on the target white anatomical surface *(TAS),* where source and target represent the left hemispheres of two different subjects aged 38±1 week PM A. Steps: a) Vertex correspondence between the source sphere *(SSS)* and anatomy *(SAS)* means that points form triplets on both surfaces; b) Control-point grids (G, red) constrain the deformation of *SSS* within a discrete optimisation scheme (orange box), c) Each control-point (blue dot) can move to a finite number of possible positions on the surface (purple crosses). The optimal displacement (blue cross) improves feature map similarity whilst constraining deformations to be smooth; d) The displaced spherical surface configuration *MSS* is estimated from ∈ using barycentric interpolation (Eq. 7); e) Barycentric correspondences are learnt between vertices on *MSS* (yellow dots) and *TSS* (pink crosses; Eq. 9); f) Weights (calculated during step e) are applied to the equivalent points on the target anatomical surface *TAS* (Eq. 10); creating g) a deformed anatomical surface configuration *(DAS),* which has the mesh topology of the source surface, but the shape of the target anatomical surface *(TAS).* Through this a transformation **F** can be estimated between *SAS* and *DAS.*

MSM employs a multi-resolution approach. In this, a sequence of spherical, regularly-sampled, control-point grids (**G**_D_)_D∈;ℕ_ are used to constrain the deformation of *MSS* (Fig. 2b). These are formed from regular subdivisions of icospheric, triangulated meshes, where the granularity of the *D*th control-point grid increases at each resolution, allowing features of the data to be matched in a coarse-to-fine fashion. Typically, at each resolution, the features from *MSS* and *FSS* are downsampled onto regular data grids *MSS_D_* and *FSS_D_* to speed processing. Resampling the data in this way may also reduce any impact that the meshing structures of MSS and FSS might have on the deformation. Final upsampling of the control-point warp to MSS is performed using barycentric interpolation.

At each resolution level, registration proceeds as a series of discrete displacement choices. At each iteration, points **p** ∈ **G**_D_, are given a finite choice of possible locations on the surface to which they can move. The end points of each displacement are determined from a set of *L* vertex points defined on a regular sampling grid (Fig. 2c, purple crosses, see also Robinson et al. (2014)). Displacements are then defined in terms of a set of *L* rotation matrices *S*^p^ = {**R^1^, R^2^.R^*L*^**}, specified separately for each control-point **p**. These rotate **p** to the sample vertex points by angles expressed relative to the centre of the sphere.

The optimal rotation for each control-point **R**_p_ ∈ *S*^p^ is found using discrete optimisation (Robinson et al., 2014; Glocker et al., 2008). This balances a unary data similarity term *c*(**R**_p_) with a pairwise penalty term *V*(**R_p_, R_q_**), which encourages a smooth warp. The search for the optimal rotations can be defined as cost function (*C*) over all unary and pairwise terms as:

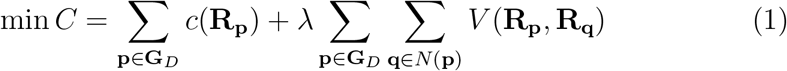

Here **q** ∈ *N*(**p**) represents all control-points that are neighbours of **p**, and *λ* is a weighting term that balances the trade off between accuracy and smoothness of the warp.

One advantage of the discrete framework is that cost functions do not need to be differentiable, and thus there are no constraints on the choice of data similarity term *c*(**R**_p_), for example: correlation, Normalised Mutual Information (NMI), Sum of Square Differences (SSD), and alpha-Mutual Information (α-MI, useful for multimodal alignments) (Neemuchwala, 2005; Robinson et al., 2014) can all be applied.

In this original framework, smoothness constraints are imposed through pairwise regularisation terms, implemented by penalising differences between proposed rotation matrices for neighbouring control-points:

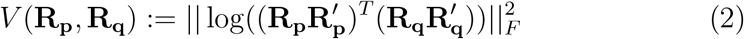

Here ||.||_*F*_ represents the Frobenius norm, and 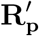 represents the full rotation of the control-point **p**, accumulated over previous labelling iterations. In what follows we refer to this regularisation framework as *sMSM_PAIR_*.

## 3. Higher-order Smoothness Constraints and Strain-based Regularization

A limitation of the original discrete optimisation framework (Robinson et al., 2014) has been that it limited regularisation and matching to first-order terms. In this paper, we compute both similarity and regularisation using whole triangles, rather than pairs of vertices, to obtain smoother, more accurate solutions. To accomplish this in a discrete framework, we apply recent advances in discrete optimisation (Ishikawa, 2009, 2014) that allow adoption of higher-order smoothness constraints.

This necessitates generalisation of the original cost function (equation 1) to allow for terms (formally known as cliques) to vary in size:

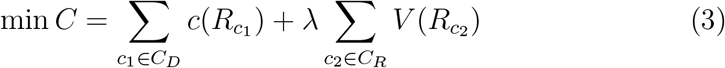

Here *C_D_* represents cliques used for estimation of the data similarity term, *C*_*R*_ represents regularisation cliques, and *R_c1_, R_c2_* represent the subset of rotations estimated for each clique. This represents a highly modular framework where any combination of similarity metric and smoothness penalty can be used, provided they can be discretised as a sum over clusters of nodes in the graph. In this paper we focus on two new triplet terms:

- **Triplet Regularisation *V(R_c2_)***: We propose a new regularisation term derived from biomechanical models of tissue deformation. This term is inspired by the strain energy minimization approach used in Knutsen et al. (2010), which constrains the strain energy density (*W_pqr_*) of locally affine warps F*_pqr_*, defined between the control-point mesh faces 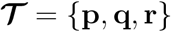. Specifically, **F***_pqr_* represents the 2D transformation matrix, or deformation gradient, for 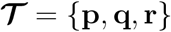 after projection into the tangent plane, fully describing deformations on the surface. The eigenvalues of **F** represent principal in-plane stretches, λ_1_ and λ_2_, such that relative change in area may be described by *J* = λ_1_λ_2_ and relative change in shape (aspect ratio) may be described by *R* = λ_1_/λ_2_. Areal distortion is traditionally defined as log_2_(*Area_1_/Area_2_*) = log_2_J. Here we introduce a similar term, “shape distortion”, defined as log_2_R. To penalize against both types of deformations, we define strain energy density using the following form:

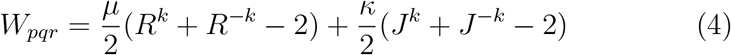

where *k* is defined as any integer greater than or equal to 1. This proposed form meets the criteria of a hyperelastic material (a class often used to characterize biological soft tissues including brain). As described for other hyperelastic materials, shear modulus, *µ*, penalizes changes in shape, while bulk modulus, *κ*, penalises changes in size (volume for traditional 3D forms, area for 2D forms). Our chosen form ensures that expansion (*J* > 1) and shrinkage (*J* < 1) are penalised equally in log space (doubling area is penalised the same as shrinking area by half). Furthermore, changes in shape (*R*) or area (*J*) are penalised by the same function to allow optimization of the trade off between areal distortion and shape distortion. Note, for the case of *k* = 1, it can be shown that this form is equivalent to a modified, compressible Neo hookean material, similar to the original form used in Knutsen et al. (2010, 2012). See Supplementary Material for more details and formal justification of the proposed strain energy form. The strain energy penalty is implemented as:

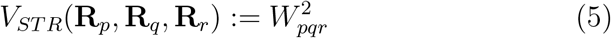
- **Triplet Likelihood *V(R_c2_)***: First proposed for registration in Glocker et al. (2010), triplet likelihoods were introduced as a means of setting up a well-posed image matching problem, since (for spherical registration) a two-dimensional displacement must be recovered from a onedimensional similarity function. As in Glocker et al. (2010) we implement triplet likelihood terms as correlations (*CC*) between patches of data: defined as all data points which overlap with each control-point mesh face triplet:

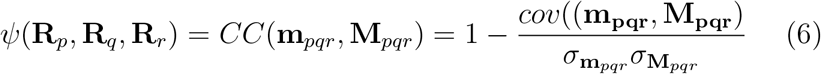 Here, **m_pqr_** is the sub-matrix of features from **m**, which correspond to points from the moving mesh *MSS* that move with the control-point triplet 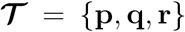. **M*_pqr_*** represent the overlapping patch in the fixed mesh space TSS. Features **M*_pqr_*** are resampled onto MSS using adaptive barycentric resampling (Glasser et al., 2013), and σ_mpqr_ and σ_Mpqr_ represent the variances of each patch.

## 4. Anatomical Regularisation (aMSM)

### 4.1. Inferring Anatomical Correspondences

Using the known vertex correspondence between the fixed sphere (*TSS*) and its anatomical representation (*TAS*, Fig. 2e) a deformed anatomical surface configuration (*DAS*) can be found for the moving surface in the following steps:

1. The coordinates (x’) of the moving source sphere (*MSS*) are found by interpolating coordinates from the control-point grid triplet 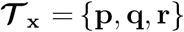 that overlaps with x, such that:

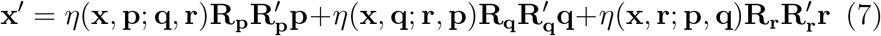 Here, 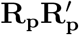 represents the combined rotation of the control-point over all iterations, and *η*(·) is a barycentric interpolation function:

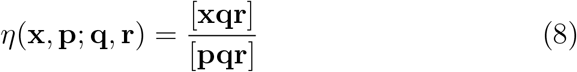

where […] represents triangle area.
2. Barycentric correspondences are found between the moving spherical surface configuration (*MSS*) and the fixed sphere (*TSS*). These are used to define a set of vertex indices, and corresponding weights, sufficient for resampling coordinates from the fixed mesh topology onto the moving mesh topology

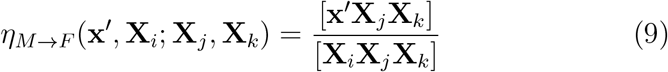 For 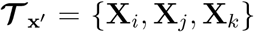, where 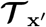 is a triplet of points on the fixed surface (FSS) that overlaps x'.
3. The indices and weights found in Eq. 9 are used to project the moving surface mesh topology onto the fixed anatomical surface. We call this the deformed anatomical surface (*DAS*) as this implements the warp that is implied through the allocation of point-wise correspondences during the spherical warp:

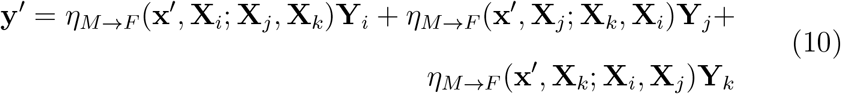

### 4.2. Implementing aMSM Regularisation

Using the higher-order constraints formulated in section 3, the evolution of DAS can be controlled by replacing spherical control-point triplets, with anatomical triplets in Equation 4. In the simplest case this can be achieved by introducing low-resolution anatomical surfaces at the resolution of the control-point grid *G_D_*. In this way a low-resolution deformed anatomical configuration *DAS_D_* can be determined from correspondences found between the control-point grid and a fixed low-resolution target sphere, *TSS_D_*, and associated low-resolution anatomical grid *TAS_D_*, with coordinates **y**, 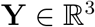 and 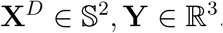. Coordinates 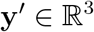 are then estimated using:

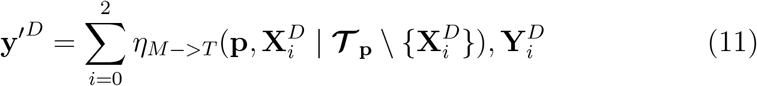

This is shorthand for the notation in Eq 10, where 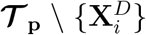 represents the points in the triplet 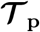 excluding 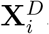. Therefore, transformations (**F** in Fig. 2) are assessed by comparing matching triplets between **y**^*D*^ and **y**^*′D*^. Nevertheless, it is important to note that the discrete displacement space continues to be estimated on the sphere. This allows registration to achieve lower anatomical distortions without sacrificing the quality of the alignment. We refer to this adaptation as anatomical MSM (*aMSM*).

## 5. Implementation details

Due to local minima in the similarity function and nonlinear penalty terms, the proposed registration cost functions are non-convex. This means that they cannot be solved by conventional discrete solvers such as *α*-expansion (used in graph cuts, Boykov et al. (2001)), which would require pairwise terms to be submodular (meet the triangle inequality). Instead, it is necessary to use methods that account for non-submodularity, such as FastPD (Komodakis and Tziritas, 2007; Komodakis et al., 2008) and QPBO (Rother et al., 2007). However, these methods only solve pairwise Markov Random Field (MRF) functions.

In order to account for triplet terms we adopt the approach of Ishikawa (2009, 2014). This allows reduction of higher order terms either by: A) addition of auxiliary variables (for example by reducing a triplet to three pairwise terms (Ishikawa, 2009); or B) by reconfiguration of the polynomial form of the MRF energy, until the higher-order function can be replaced by a single quadratic (Ishikawa, 2014). We present results using the latter version, known as Excludable Local Configuration (ELC). This has the advantage that, provided an ELC can be found, there is no increase in the number of pairwise terms, which has some impact on the computational time.

Once terms are reduced, optimisation proceeds as solutions to a series of binary label problems, where results for the full label space are obtained using the hierarchical implementation of the fusion moves technique (Lempitsky et al., 2010), as described in Glocker et al. (2010). In each instance, the reduction is passed to the FastPD solver (Komodakis and Tziritas, 2007; Komodakis et al., 2008) for optimisation.

For effective *aMSM* implementation, choices have to be made with regards to how far the anatomy should be reasonably downsampled. Resampling of the anatomical surface is performed by barycentric interpolation, using correspondences between the initial source sphere (*SSS*) and regular, low-resolution icospheric spherical grids (Supplementary Material). Where resolution of *SAS^D^* exceeds that of the control-point grid, regularisation for each control-point triplet is calculated by averaging distortions for every higher-resolution triplet in *SAS^D^* that falls within the area of that control-point grid face. This results in a trade-off between run time and accurate capture of the true anatomical distortion (see Supplementary Material). For every increase in anatomical mesh resolution relative to the control-point grid, strain calculations better represent that of the native deformation, but number of strain calculations increases by a factor of four.

In general, estimation of triplet energies and reduction through ELC and binary FastPD slows the run time relative to the original MSM approach. To reduce some of the impact, the control-point grids are no longer reset after each iteration. Instead, the source mesh and control grid incrementally deform together and all neighbourhood relationships are learned once at the beginning of each resolution level. To prevent folding of the mesh during alignment a weighting penalty is placed on the regularisation cost that severely penalises flipping of the triangular faces.This is done to ensure the final transformation is smooth and invertible.

Finally, to improve convergence and allow for smoother warps, modifications are also made to the rescaling of the discrete label space at each iteration. In the original framework the label space switches between the vertices and barycentres of a regular sampling grid (Robinson et al., 2014). In the new framework the discrete displacement vectors 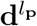 are instead rescaled by 0.8× their original length over 5 iterations, before being reset to their original lengths.

## 6. Experimental Methods and Results

We tested this framework on real data collected as part of the adult Human Connectome Project (HCP^4^) and Developing Human Connectome Project (dHCP^5^) to assess the impact of the proposed strain regulariser on both spherical and anatomical deformations. In all experiments results were compared for the higher-order MSM registration framework (*MSM_STR_*) and the original form (*sMSM_PAIR_*). Where feasible MSM was also compared against FreeSurfer (*FS*: arguably the most commonly used tool for cortical surface alignment) and Spherical Demons (*SD*), which is diffeomorphic on the sphere. Note, standard implementations of FreeSurfer and Spherical Demons are not capable of multimodal or multivariate alignment.

### 6.1. Cohorts

A subset of 28 subjects for the full HCP cohort (1200 subjects) were selected for parameter optimisation of the multimodal alignment protocol laid out in Glasser et al. (2016a). Here registration was driven using a combination of features reflecting myelin maps (Glasser and Van Essen, 2011), 34 well-defined cortical surface resting-state fMRI spatial maps, and 8 visuo-topic features (reflecting topographic organisation of functional connectivity in the visual cortex; see Glasser et al. (2016a) Supplementary Methods sec 2). This subset of subjects was selected to reflect a spectrum of sources of variance in the HCP data, including data sets with unusual functional topology (see Glasser et al. (2016a) Supplementary Results sec 1.3); or low signal-to-noise ratio (SNR).

In the second experiment a subset of dHCP subjects were used to explore longitudinal alignment of developing cortical shapes. This group was selected specifically to include all subjects scanned twice within 32.66 ± 1.22 weeks PMA (first scan) and 41.47 ± 1.61 weeks PMA (second scan) in order to allow straight-forward comparison of the deformations across subjects.

### 6.2. Data

Acquisition of HCP data was performed on a Siemens 3T Skyra platform, using a 32-channel head coil and MPRAGE (T1w) and SPACE (T2w) sequences (Glasser et al., 2013). Isotropic structural image acquisitions were acquired at 0.7mm^3^. Functional imaging data was acquired with multiband (factor 8) 2mm Gradient-Echo EPI sequences (Moeller et al., 2010). Four resting state fMRI (rfMRI) scans were acquired (two successive 15 minutes scans in each of two sessions) (Smith et al., 2013). Seven task-fMRI (tfMRI) experiments were also conducted, including: working memory, gambling, motor, language, social cognition, relational, and emotional tasks; tfMRI scans were acquired after the rfMRI scans in each of two hour-long sessions on separate days (Barch et al., 2013). Additional details regarding specific acquisition parameters and task protocols are available in Barch et al. (2013) and Smith et al. (2013), as well as on the HCP website^6^. Generation of surface meshes and associated shape features were carried out using HCP Structural Pipelines (Glasser et al., 2013).

dHCP data was acquired at St. Thomas Hospital, London, on a Philips 3T scanner using a 32 channel dedicated neonatal head coil (Hughes et al., 2016). To reduce the effects of motion, T2 images were obtained using a Turbo Spin Echo (TSE) sequence, acquired in two stacks of 2D slices (in sagittal and axial planes), using parameters: TR=12s, TE=156ms, SENSE factor 2.11 (axial) and 2.58 (sagittal). Overlapping slices (resolution (mm) 0.8×0.8×1.6) were acquired to give final image resolution voxels 0.8 × 0.8 × 0.8mm^3^ after motion corrected reconstruction, combining Cordero-Grande et al. (2016); Kuklisova-Murgasova et al. (2012). T1 images were acquired using an IR-TSE (Inversion Recovery Turbo Spin Echo) sequence at the same resolutions with TR=4.8s, TE=8.7ms, SENSE factor 2.26 (axial) and 2.66 (sagittal). All images were reviewed by an expert paediatric neuroradiologist and checked for possible abnormalities. Generation of surface meshes and associated shape features were carried out using dHCP Structural Pipelines (Makropoulos et al., 2017).

### 6.3. Strain Parameter Optimisation

Higher-order MSM (MSM_STR_) was run using: tri-clique data terms, bulk modulus (*κ*) of 1.6, shear modulus (*µ*) of 0.4 (i.e. a 4 to 1 ratio), and *k* = 2. These parameters were empirically optimised for multimodal alignment of adult HCP data (sec. 6.4), keeping *κ* + *µ* = 2 and regularisation *λ* constant, for maximum alignment and minimum total deformations. These values were kept throughout the paper. Experiments on the influence of these parameters for longitudinal alignment of cortical anatomies (run using *aMSM*_*STR*_, Supplementary Material sec. 2) show that, in general, results are robust over a range of parameters.

### 6.4. Multimodal alignment of adult HCP data

In the first experiment, *sMSM_STR_* was compared against *sMSM_PAIR_*, *SD*, and *FS* for alignment of task fMRI data from a subset of 28 subjects from the HCP project. In this instance, anatomical regularisation was not used lest it predjudiced the solution towards alignment of cortical folding patterns, as these may not consistently reflect areal features across large parts of the brain across subjects. MSM methods were compared against *FS* and *SD*, run using their default settings (cortical folding alignment only) because *FS* registration is fixed and immutable, and the current implementation of *SD* allows only for alignment of univariate features.

MSM was run in two stages: first registration was initialised using constrained alignment of cortical folds (as described in Robinson et al. (2014) as the MSMSulc procotol) optimized in order to maximize task fMRI alignment. Then alignment of areal features was refined using what has become known as MSMAll (Glasser et al., 2016a). In this multimodal registration features containing myelin maps (Glasser and Van Essen, 2011), resting-state networks (RSNs) and visuotopic features were used to drive alignment to a group average template.

MSMSulc was run using strain-based regularisation (*sMSM_STR_*) over three control-point (*CP*) grid resolutions with *CP* resolution: *CP_res_*=162, 642, 2542; and features sampled to regularly-spaced grids (*MSS_D_,FSS_D_*) at resolutions: *DP_res_*=2542, 10242, 40962. Regularisation strength was controlled through λ = 10, 7.5, 7.5. Features were variance normalised but no smoothing of the data was performed. Notably, the re-optimised strain-based MSMSulc substantially outperforms both pairwise MSMSulc and the FreeSurfer registration (Table 1).

MSMAll was optimised three times, once using strain-based higher-order regularisation and likelihood terms (sMSM_STR_) and twice using pair-wise regularisation *sMSM_PAIR_*. In each case, common parameters between the methods were fixed; registration was run over three control-point grid resolutions: *CP_res_*=162, 642, 2542, *DP_res_*=2542, 10242, 40962; features were variance normalised and there was no smoothing of the data. Regularisation of *sMSM_STR_* was optimised in order to maximise alignment of tfMRI (appraised through estimates of cluster mass - defined below), subject to mean edge distortion (defined below) not exceeding that for FreeSurfer. Then *sMSM_PAIR_* was optimised twice: in the first case to achieve comparable peak edge distortions to *sMSM_STR_* (this keeps peak distortions well controlled, but limits the amount of registration that can occur, see introduction), and in the second case to match comparable mean values of edge distortions (this allows a similar amount of registration to occur, but peak distortions may exceed neurobiologically plausible thresholds).

Following registration, tfMRI timeseries on the subjects’ native meshes were resampled according to the registration to the standard mesh. Improvements in alignment were assessed via comparisons of the group mean activation maps (obtained using mixed effects FLAME Woolrich et al. (2009)) both qualitatively and quantitatively, via cluster mass, calculated using the following formula: 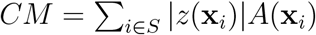. Here *x_i_*, is a vertex coordinate, *z*(x*_i_*) is the statistical value at this coordinate, *A*(x*_i_*) is the area associated with this vertex (calculated from a share of the area of each mesh triangle connected to it in the mid-thickness surface), and *S* is the set of vertices where |*z*(x*_i_*| > 5, and the threshold of 5 was chosen to be approximately equivalent to a two-tailed Bonferroni correction. The cluster mass measure reflects both the size of the super-threshold clusters and the magnitude of the statistical values within them.

Distortions are reported in terms of absolute values for: areal distortions (log_2_ J, Eq. 4), shape distortions (log_2_ R), and edge distortions. Edge distortions are estimated from the relative change in length of edges between neighbouring vertices in the mesh: 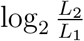, where *L_2_* is edge length following registration, and *L_1_* is length before, and are reported per vertex by taking the average values for all connected edges. These maps reflect a univariate summary of changes to area and shape and thus were used during optimisation

Results in Table 1 and Fig. 3 demonstrate that, as expected, multimodal alignment of tfMRI data (MSMAll) significantly improved the sharpness and peak values of the group z-statistics, relative to cortical folding based alignment (*SD, FS, MSM*Sulc). This resulted in a 20.97% increase in total cluster-mass for MSMAll run with *sMSM_STR_* relative to MSMSulc, a comparable increase of 22.42% over SD, and a 27.31% increase over FS. Fig. 5 displays the spread of improvements across all pure contrasts within each task as improvement relative to *FS* - again *sMSM_STR_* outperformed all other methods.

**Figure 3.**
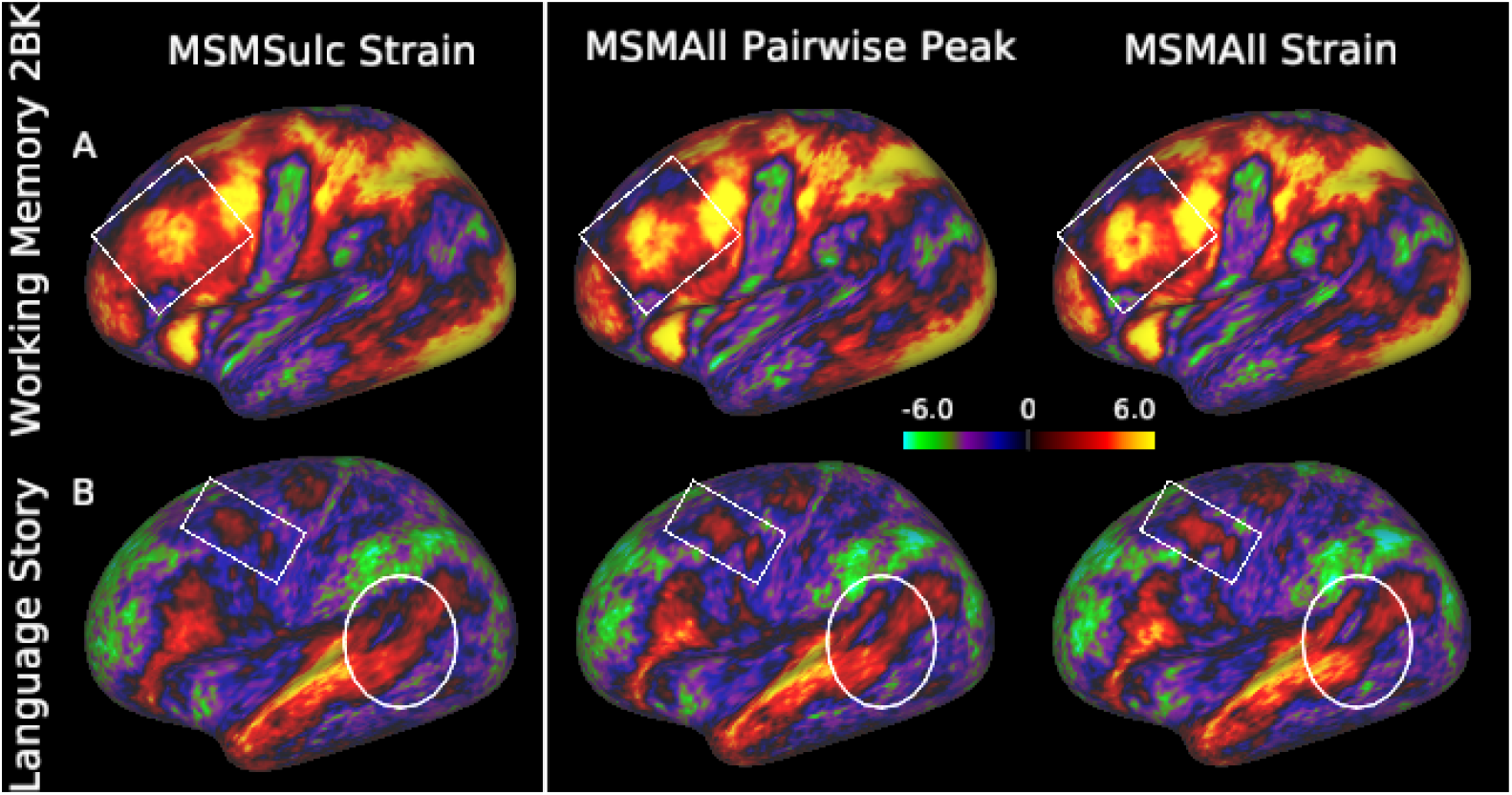
Comparison of group Z-statistic spatial maps following folding alignment (MSM-Sulc, run with *sMSM_STR_*) and alignment driven my multimodal features (MSMAll, run with *sMSM_STR_* and *sMSM_PAIR_*, matched for peak strains) for: a) a working-memory contrast (2BK) and b) a language task (Story). White boxes highlight improvements in sharpness of the contrast in the areas of the Dorsal Lateral Pre-Frontal cortex (A); region 55b (B) and in the temporal lobe (B)

In terms of folding-based methods, both *sMSM_STR_* and *SD* produced much lower and smoother distortions than *FS*; although it is important to note both were optimised for alignment of areal features (Robinson et al., 2014; Yeo et al., 2010), whereas *FS* was optimised for alignment of cortical folds. MSMSulc achieved marginal improvements over *SD*, with slight increases in tfMRI cluster mass obtained from deformations with lower isotropic distortions, and similar edge distortions. Fig. 6 A-C (top row) shows *sMSM_STR_* edge distortions dispersed across the whole of the surface whereas *SD* alignment resulted in peaks of high distortions.

For multimodal alignment, optimising the original form of MSM to achieve comparable mean distortions achieved comparable improvements in cluster mass to *sMSM_STR_* but led to extensive patches of extreme distortions across the cortical averages (Fig 4, Fig 6D), with peak values for log_2_ J (Fig 4 left) and log_2_ R (Fig 4 right) far exceeding that of *sMSM_STR_* by 128% and 63% respectively. Such levels of distortions are neurobiologically highly implausible given expected ranges of regional variation from previous studies (Van Essen, 2005).

**Figure 4.**
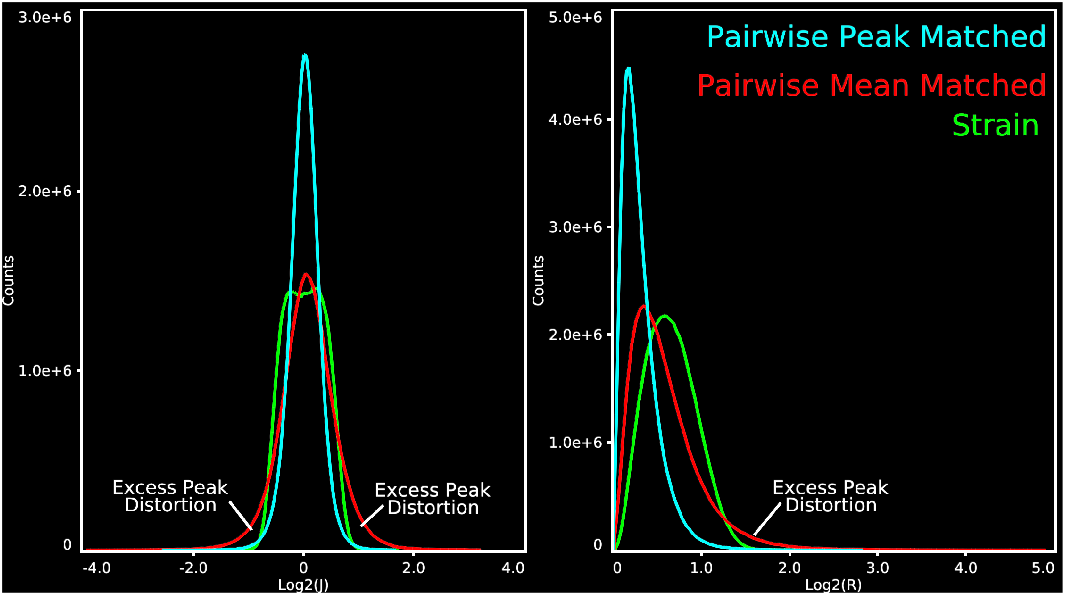
Histogram plots comparing MSMAll distortions, plotted against log_2_ J (left) and log_2_ R (right). *sMSM_PAIR_* registration generates long tailed distributions with excessive peak distortions

**Figure 5.**
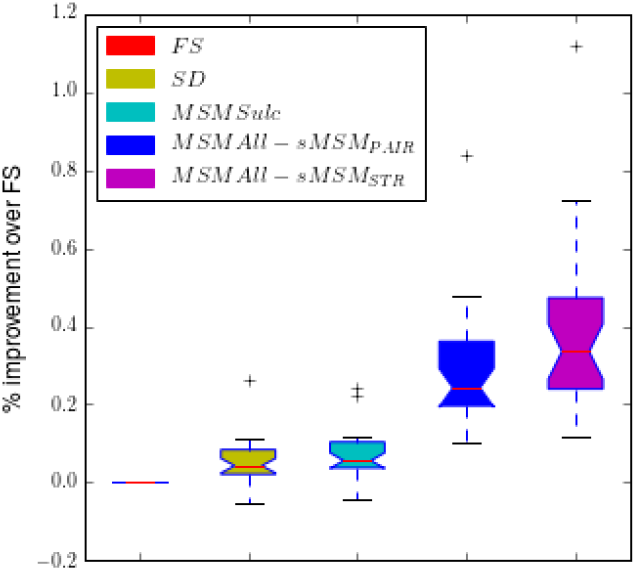
Bar chart of mean cluster mass statistics across HCP task categories for different methods. Note, only pure contrasts are included, that is direct response to individual tasks not differences in activations between tasks.

**Figure 6.**
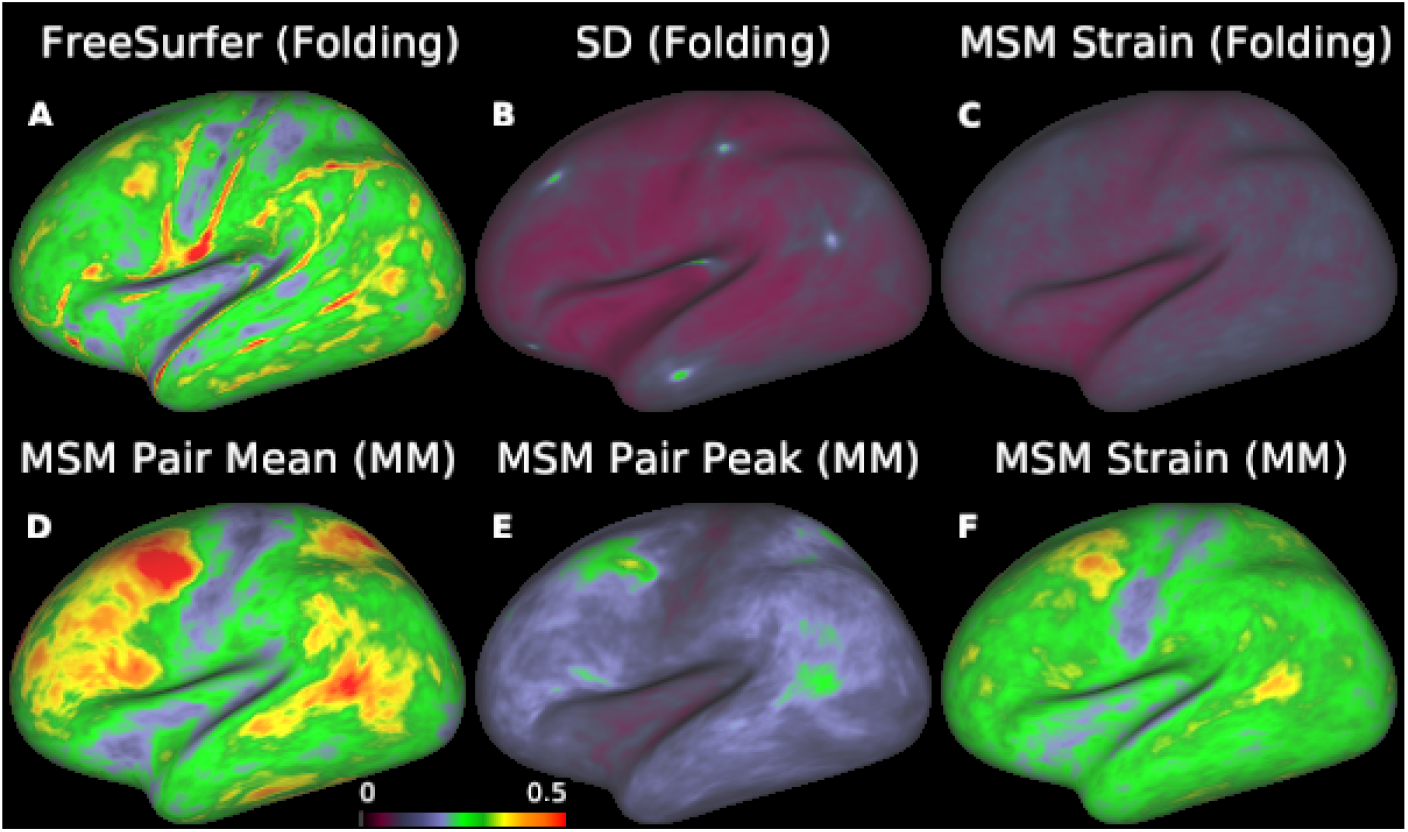
Mean edge distortion maps, averaged across all surfaces. Top row) distortions for folding based alignment only, pink boxes highlight hot spots of edge distortions for *SD* method; Bottom row) multimodal (MM) alignments: MSM Pair Mean (MSMAll run with sMSMpAiR optimised to achieve comparable mean strains to *sMSM_STR_*); MSM Pair Peak (MSMAll run with *sMSM_PAIR_* optimised to achieve comparable peak strains to *sMSM_STR_*; MSM Strain (MSMAll run with *sMSM_STR_*)

When distortions were instead matched for peak strains, improvements in total cluster mass observed for *sMSM_STR_* were 3.86% above the gains obtained with the original form of MSM (Table 1). This resulted in sharper tMRI group task maps (Fig. 3). Note, greater improvements would be expected should regularisation of *sMSM_STR_* be reduced further. However, out of an abundance of conservatism we chose not to exceed the edge distortion of FreeSurfer.

**Table 1.** Peak distortions and cluster mass (mm^2^) estimates following alignment of HCP tfMRI data using each of the proposed methods.

|  | Cluster Mass | Edge distortion |  | Areal distortion |  | Shape distortion |  |
| --- | --- | --- | --- | --- | --- | --- | --- |
|  |  | Mean | Max | Mean | Max | Mean | Max |
| *FS* | 3212876 | 0.271 | 2.564 | 0.377 | 4.879 | 0.698 | 6.752 |
| *SD* | 3341178 | 0.074 | 0.520 | 0.124 | 1.087 | 0.147 | 0.884 |
| MSMSulc ($sMSM_{PAIR}$) | 3019301 | 0.089 | 0.486 | 0.142 | 0.879 | 0.183 | 1.040 |
| MSMSulc ($sMSM_{STR}$) | 3381148 | 0.089 | 0.268 | 0.102 | 0.525 | 0.235 | 1.151 |
| MSMAll ($sMSM_{PAIR}$, peak matched) | 3938268 | 0.139 | 0.972 | 0.209 | 2.561 | 0.311 | 2.824 |
| MSMAll ($sMSM_{PAIR}$, mean matched) | 4022388 | 0.271 | 1.918 | 0.387 | 3.939 | 0.604 | 4.901 |
| MSMAll ($sMSM_{STR}$) | **4090190** | 0.261 | 0.919 | 0.306 | 1.731 | 0.662 | 3.015 |

### 6.5. Longitudinal Registration

In the second experiment we explored the impact of anatomical warp regularisation for within-subject longitudinal alignment of 10 different neonatal subjects, each scanned twice within two specific time points (TPs): 32.66 ± 1.22 weeks PMA (TP1) and 41.47 ± 1.61 weeks PMA (TP2). This criterion was selected by clustering all longitudinally scanned subjects (37 at time of writing) into groups with similar TPs, such that the resulting deformations may be directly compared.

Here we present results from the group with the biggest scan separation. Over this time period cortical geometry and the pattern of distortions resulting from the spherical projection change dramatically (Fig. 7 a). However. since the relationship between structural and functional organisation of the brain presumably remains reasonably consistent, we assume it is sufficient to drive registration using geometric features only (mean curvature, Fig. 7 c).

**Figure 7.**
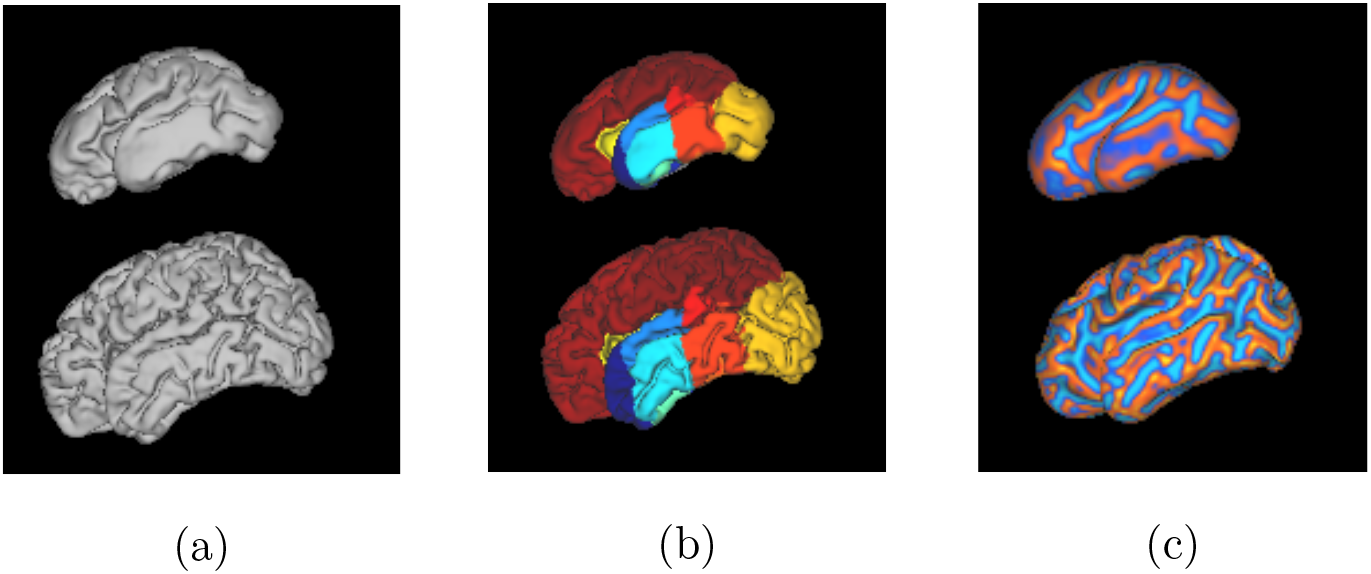
Comparison of surface geometry (a), cortical labels (b) and curvature maps (c) for one exemplar data set (top=TP1, bottom=TP2)

Results are compared for: the original MSM spherical framework (*sMSM_PAIR_*); spherical MSM with higher-order strain regularisation (*sMSM_STR_*); anatomical MSM with strain regularisation (*aMSM_STR_*); and Spherical Demons (SD). All registrations were run using the TP1 surface as the target of the registration, and TP2 as the moving surface, as this was found to generate the most accurate alignment, where this was judged in terms of correlation between features sets, relative to total surface distortion. This is likely because it is a better-defined problem to register a more complex surface to a simpler surface than the reverse. Deformations in the direction TP1 → TP2 were then obtained by inverting the transformation. Note that biases associated with unidirectional registration can be circumvented by registering in both directions and averaging, as performed in Garcia et al. (under review, bioRxiv reference: 185389).

*SD* was run using its default parameterisation and represents a baseline for smooth diffeomorphic spherical alignment. All MSM registrations were optimised over 4 resolution levels with mesh resolutions: *CP*_res_=162, 642, 2542, 10242, *DP*_res_=10242, 10242, 40962, 40962; variance normalisation and smoothing was applied σ_in_ = σ_ref_ = 6, 4, 2,1. *aMSM_STR_* was parameterised to penalise distortions of the midthickness surfaces, using anatomical mesh resolutions (*DAS_D_, TAS_D_*) of *AG_res_*=2542, 10242, 40962, 40962. To encourage even penalization of size changes across the surface, *SAS* and *TAS* were normalised to the same total surface area prior to *aMSM_STR_* registration. In each case λ was selected to optimise feature map correlation relative to areal distortions.

After registrations, true midthickness surfaces were projected through the estimated warp using the procedure described in sec. 4. Methods were compared in terms of the goodness of fit of the obtained alignments relative to distortions of the anatomy. In order to compare deformations between data sets, relative values for areal (log_2_ J) and shape (log_2_ R) distortions were estimated at each vertex *i* as 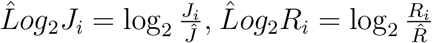, where *Ĵ* and 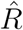 represent (per surface) average values of R and J, respectively. Alignment quality was assessed through improvements in correlation of curvature feature maps and Dice overlap of 16 folding-based cortical regions (Fig. 9) relative to affine registration. Here, cortical labels were obtained through registration of each timepoint’s T2 brain volume to 20 manually annotated neonatal atlases (ALBERTS: Gousias et al. (2012)). The resulting 20 segmentations were then fused in a locally-weighted scheme to form the subject’s cortical labels (see LWV-MSD in Artaechevarria et al. (2009); Makropoulos et al. (2014)), so that similar patches between each atlas and the image have increased weighting.

Results in Fig. 8 show strong improvements for *aMSM_STR_* over the purely spherical methods. Areal distortions are much reduced (Fig 8 a), and shape distortions (Fig 8 b) are lower than all methods other than for SD, for which alignment quality is comparatively lower (Fig. 9).

**Figure 8.**
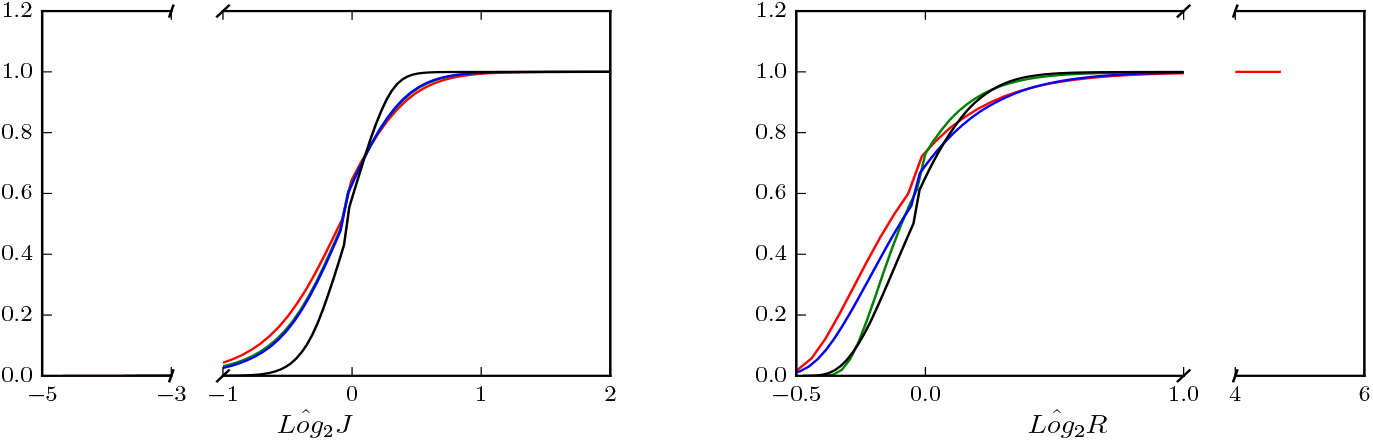
Cumulative distribution functions of *L̂og*_2_*J* and *L̂og*_2_*R*. for different methods: *sMSM_PAIR_* (red); *SD* (green); *sMSM_STR_* (blue); *aMSM_STR_* (black). Functions are estimated from the full distribution of strain values estimated by combining per-vertex strain values across all 10 deformations.

**Figure 9.**
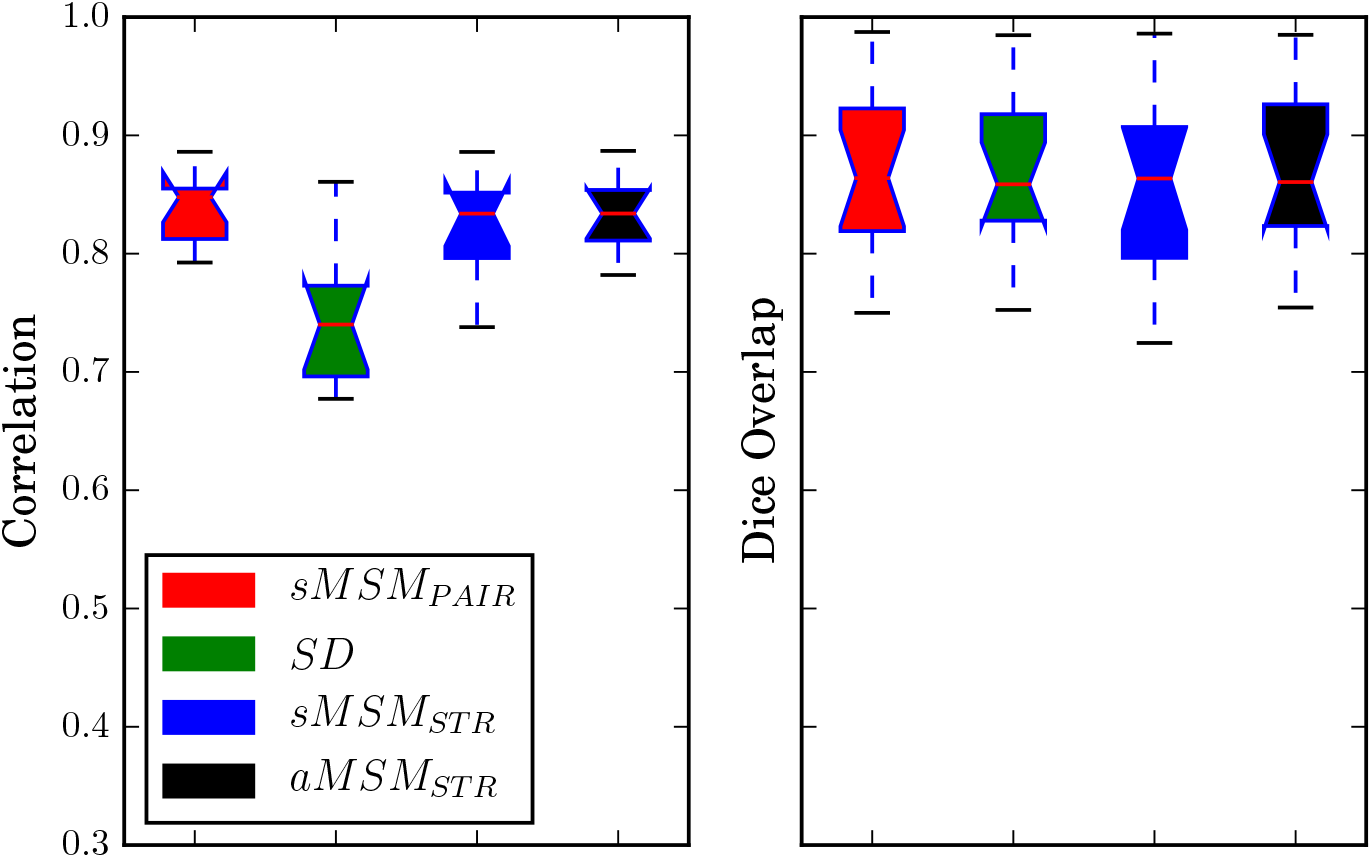
Alignment quality of longitudinal warps, assessed through feature map cross correlation and Dice Overlap (averaged across 16 cortical regions). Colour as for Fig. 8

To further assess the smoothness and consistency of the distortions between subjects, the initial time point for all scans was registered to a 34 week surface template (Bo΂ek et al., 2016), using *sMSM_STR_* alignment of sulcal depth maps using the following parameters: *CP*_res_=162, 642, 2542; *DP*_res_=2542, 10242, 40962; λ = 0.5, 0.5, 0.5; σ_in_ = σ_ref_ = 6, 4, 2. Distributions of *LogJ* for each subject were then resampled onto the template and compared using FSL’s PALM (Winkler et al., 2014), which performs permutation testing for surface image data. This assesses at each vertex whether distortions are statistically greater than zero, performing family-wise error correction through Threshold Free Cluster enhancement (TFCE,Smith and Nichols (2009)).

Results in Fig. 10 show mean *Log*_2_*J* across the surface is much smoother for *aMSM*_*STR*_ than for spherical methods (*SD*, *sMSM_PAIR_*, *sMSM_STR_*). This translates to much broader areas where distortions (expansion at TP2 vs TP1) are significantly above zero. These areas correspond to regions in the frontal and parietal lobe, which are expected to grow faster during this time period as well as after birth (Moeskops et al., 2015; Hill et al., 2010).

**Figure 10.**
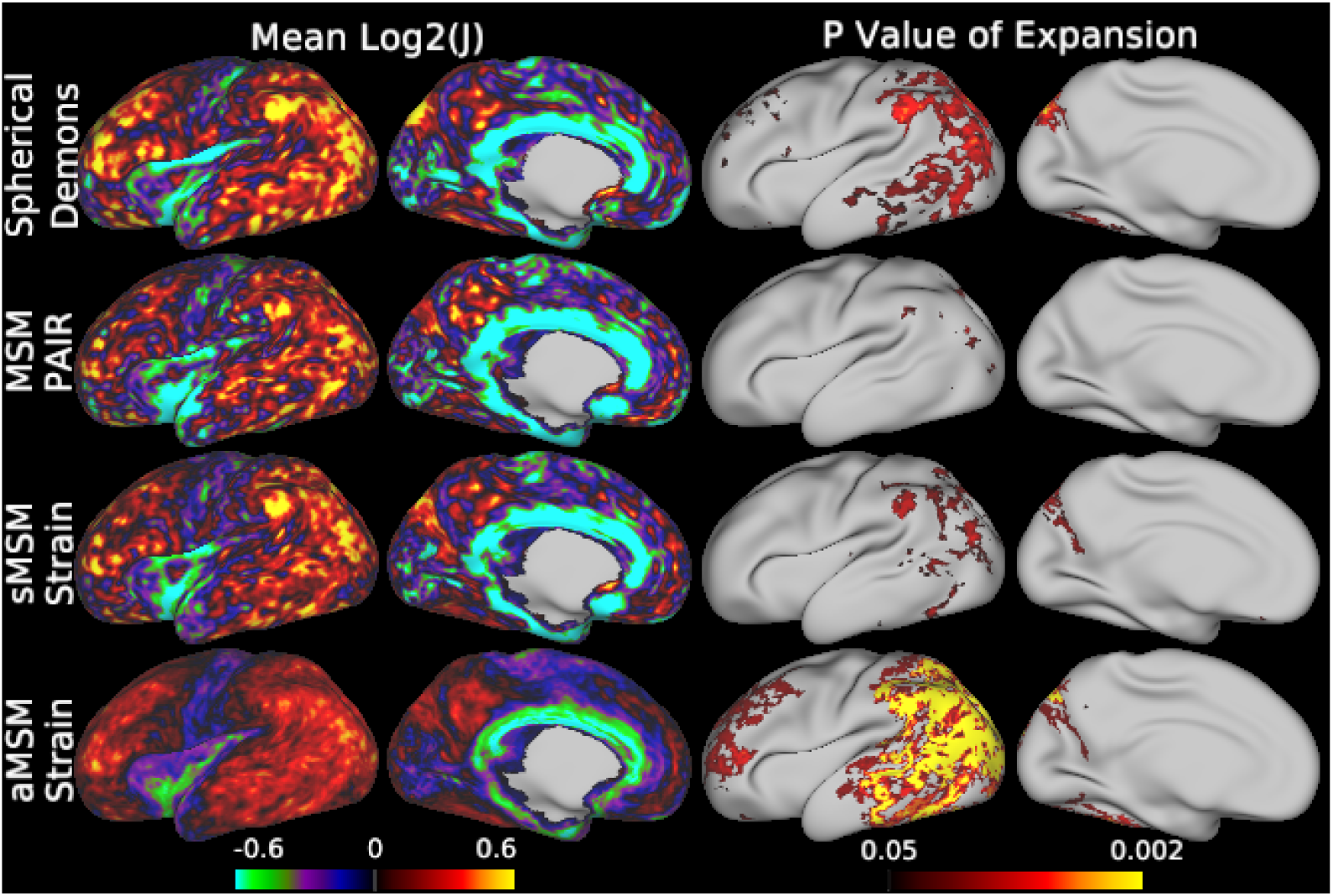
Comparison of distortion fields across subjects. Left: *L̂og*_2_*J* relative areal distortion averaged across all 10 neonatal subjects in template space; Right: pvalues for statistical comparison, thresholded at *p* < 0.05

## 7. Discussion

Human brain imaging studies are extremely diverse; a wide variety of different tissue properties are studied using a range of available imaging modalities. The relationships between these properties are unknown but are likely to be complex since studies have already shown disassociations between patterns of functional and folding organisation (Amunts et al., 2007; Glasser et al., 2016a). Despite this issue the majority of spatial normalisation techniques focus on alignment of specific feature types, including: cortical folding patterns (Fischl et al., 1999b; Yeo et al., 2010; Wright et al., 2015; Lombaert et al., 2013), cortical geometry (Durrleman et al., 2009), tfMRI time series (Sabuncu et al., 2010), patterns of functional organisation (Conroy et al., 2013; Nenning et al., 2017), and fixed combinations of geometry and fMRI/dMRI (Lombaert et al., 2015; Orasanu et al., 2016b). By contrast in this paper we present a method which allows flexible and robust alignment of a wide variety of different combinations of features on the cortical surface, as well as biologically-constrained and statistically-plausible deformations of cortical anatomy.

This paper builds on previous work in which we proposed a tool for spherical alignment of brain imaging data, implemented using discrete optimisation (Robinson et al., 2014). This framework offered significant advantages for multimodal registration on account of offering a modular and flexible choice of cost functions, and deformations robust to local minima. One limitation of the original approach was that it was implemented through use of first-order discrete methods and a regularization function with several limitations (see introduction). This created tension between controlling peak distortions in regions with individual differences in the layout of cortical areas, and maximizing alignment across the rest of the surface. This paper therefore presents an improved framework that utilises advances in discrete optimisation to include higher-order smoothness penalty terms (Ishikawa, 2009, 2014). Using this framework we improve the robustness of MSM through inclusion of a new hyperelastic strain energy density penalty (Eq. 4) that allows complete control over local changes in shape (*R*) and area (*J*). In this way we have been able to significantly improve the alignment of complex multimodal feature sets. Notably, this includes, enhancing the alignment of data from the adult multimodal parcellation feature set from the HCP (Glasser et al., 2016a).

A further advantage of the proposed higher-order framework is that it enabled regularisation of the *anatomical* deformations implied by the spherical warp. Experiments performed for longitudinal alignment (10 neonatal cortices) show that anatomical MSM (*aMSM*) generates smooth and biologically plausible deformations. These deformations reflect patterns of cortical growth similar to those previously reported in region-of-interest studies (Moeskops et al., 2015). But, importantly, *aMSM_STR_* allowed us to observe statistically significant trends using a much smaller sample size, and without the loss of detail associated with large regions of interest.

Throughout this paper we have compared the proposed MSM only against other spherical registration frameworks: Spherical Demons (*SD*) and FreeSurfer (*FS*), and solely optimised for cortical folding alignments. We compare directly to spherical methods as these offer more flexibility in terms of the range of features that can be used to drive the registration. Whilst it is possible that methods designed for smooth and/or diffeomorphic alignment of cortical surface geometries (Durrleman et al., 2009; Lombaert et al., 2013; Orasanu et al., 2016a) may outperform MSM for the specific task of longitudinal alignment, to our knowledge neither have reported smooth, statistically significant surface expansion maps like those produced by aMSM. Further, these methods are coupled to alignment of cortical shape. This limits their flexibility for between subject alignment on account of known dissociation between the brain’s functional (or areal) organisation and patterns of cortical folding (Amunts et al., 2007; Glasser et al., 2016a; Nenning et al., 2017). For the same reason, it is not clear how methods that combine spectral alignment of shape with correspondences learnt from functional or diffusion MRI (Lombaert et al., 2015; Orasanu et al., 2016a) should resolve this conflict to obtain a unified mapping across the whole brain.

Here we compare against *SD* and *FS* only for folding alignment on account of the fact that *FS* registration is fixed and hard-coded (allowing cortical folding alignment only) and the current implementation of *SD* allows only for alignment of univariate features. It is true that other groups have applied *SD* to alignment of brain function by mapping or embedding functional data to a univariate space (Nenning et al., 2017; Tong et al., 2017). However, our comparisons of *SD* and *sMSM_STR_* for registration of adult folding data show that *sMSM_STR_* achieves warps with comparable smoothness to SD. By contrast, a strength of MSM is that it works flexibly with a range of multivariate and multimodal features. Furthermore, this flexibility has overcome a limitation of traditional spherical methods by generating plausible deformations of cortical surface anatomies. For these reasons we consider an extensive comparison between MSM and these specialised cases outside of the scope of this paper, but would welcome an independent analysis of these issues.

There remain limitations of the proposed approach inasmuch as the current implementation, using Ishikawa (2009, 2014), reduces the higher-order problem to a series of binary problems. In this paper, a multi-label solution is obtained through use of the fusion moves technique (Lempitsky et al., 2010). However, this circumvents the fast multi-label optimisation offered by the FastPD algorithm (Komodakis and Tziritas, 2007; Komodakis et al., 2008), leading to a slower solution. For comparison, on a specific 64-bit Linux system, the proposed version of MSM runs in approximately 1hr 15 minutes (comparable in run time to FreeSurfer) whereas the original pairwise form runs in less than 10mins, and Spherical Demons runs in less than 5 mins. Run times can be considerably brought down through appropriate code parallelisation. However, future work should also explore alternative higher-order optimisation strategies such as Fix et al. (2014); Komodakis and Paragios (2009).

Despite improved robustness of the proposed method to the effects of noise and topological variance in the data, this method is still a spatially-smooth, topology-constraining registration approach. An important future avenue will be to address the significant limitation that spatially constrained deformations cannot align brains with variable functional topologies (such as those observed for in ~10% of subjects for area 55b in Glasser et al. (2016a)). One avenue may be to explore combining hyper-alignment (Langs et al., 2010; Haxby et al., 2011), or graph matching (Ktena et al., 2016), approaches with spatially-constrained registration, such that constraints are placed to ensure that regions cannot be matched if they are very far apart in space (Iordan et al., 2016). Alternatively, in Robinson et al. (2016) we propose a group-wise registration scheme that accounts for topological variation though minimisation of rank of the feature set across the group.

In conclusion, we believe that this study establishes MSM as a powerful and flexible tool, which provides a valuable resource for studying a wide variety of properties of brain organisation across a range of populations. The strain-based version of MSM will be used in HCP studies on development, aging, and disease. Future work will expand studies on longitudinal fetal and neonatal cortical development across a larger cohort to better understand the mechanisms underpinning cortical growth, and will extend studies of neonatal resting-state networks through development of spatio-temporal templates of brain functional and structural organisation Bo΂ek et al. (2016). By quantifying patterns of structural and functional development, it may be possible to generate vital biomarkers indicating neurodevelopmental outcomes for vulnerable groups such as preterm infants.

## 8. Acknowledgements

The research leading to these results has received funding from the European Research Council under the European Unions Seventh Framework Programme (FP/2007-2013) / ERC Grant Agreement no. 319456. We are grateful to the families who generously supported this trial. The work was supported by the NIHR Biomedical Research Centers at Guys and St Thomas NHS Trust

Data from the Human Connectome Project (HCP - WU-Minn Consor-tium, Principal Investigators: David Van Essen and Kamil Ugurbil; 1U54MH091657 was funded by the 16 NIH Institutes and Centers that support the NIH Blueprint for Neuroscience Research; and by the McDonnell Center for Systems Neuroscience at Washington University. MFG was supported by an individual fellowship F30-MH097312 (NIH) and DVE by NIIH R01 MH060974. KG was supported by grant T32 EB018266 (NIH). MJ was supported by the NIHR Biomedical Research Centre, Oxford.

## Footnotes

2 note that we refer to all of the in vivo MR-based estimates of myelin such as T1w/T2w or quantitative T1 as myelin maps in this study. For further discussion, see Glasser et al 2014 Neuroimage, Glasser et al 2016a, and Glasser et al 2016b)

3 note, that the released HCP MSMAll data used the ADP version of MSMAll, however the MSMSulc and MSMAll pipelines based on strain-based regularisation are publicly available (https://github.com/Washington-University/Pipelines) and will be used in follow up HCP projects on development, ageing and human disease;

4 http://www.humanconnectome.org/

5 http://www.developingconnectome.org/

6 http://www.humanconnectome.org/documentation/Q1/imaging-protocols.html

